# Safety and Immunogenicity Evaluation of Inactivated whole-virus-SARS-COV-2 As Emerging Vaccine Development In Egypt

**DOI:** 10.1101/2021.03.01.433130

**Authors:** Amani A. Saleh, Mohamed A. Saad, Islam Ryan, Magdy Amin, Mohamed I. Shindy, Wael A. Hassan, Mahmoud Samir, Ayman A. Khattab, Sherein S. Abdelgayed, Mohamed G. Seadawy, Hossam M. Fahmy, Khaled Amer

## Abstract

The current worldwide pandemic COVID-19 is causing severe human health problems, with high numbers of mortality rates and huge economic burdens that require an urgent demand for safe, and effective and vaccine development. Our study was the first trail to development and evaluation of safety and immune response to inactivated whole SARS-COV-2 virus vaccine adjuvanted with aluminium hydroxide. We used characterized SARS-COV-2 strain, severe acute respiratory syndrome coronavirus 2 isolates **(SARS-CoV-2/human/EGY/Egy-SERVAC/2020)** with accession numbers; **MT981440**; **MT981439; MT981441; MT974071; MT974069 and MW250352** at GenBank that isolated from Egyptian patients SARS-CoV-2-positive. Development of the vaccine was carried out in a BSL - 3 facilities and the immunogenicity was determined in mice at two doses (55µg and 100µg per dose). All vaccinated mice were received a booster dose 14 days post first immunization. Our results demonstrated distinct cytopathic effect on the vero cell monolayers induced through SARS-COV-2 propagation and the viral particles were identified as Coronaviridae by transmission electron microscopy. SARS-CoV-2 was identified by RT-PCR performed on the cell culture. Immunogenicity of the developed vaccine indicated the high antigen-binding and neutralizing antibody titers, regardless the dose concentration, with excellent safety profiles.However, no deaths or clinical symptoms in mice groups. The efficacy of the inactivated vaccine formulation was tested by wild virus challenge the vaccinated mice and detection of viral replication in lung tissues. Vaccinated mice recorded complete protection from challenge infection three weeks post second dose. SARS-COV-2 replication was not observed in the lungs of mice following SARS-CoV-2 challenge, regardless of the level of serum neutralizing antibodies. This finding will support the future trials for evaluation an applicable SARS-CoV-2 vaccine candidate.

## Introduction

Coronavirus disease 2019 (COVID-19) is an emerging respiratory infectious disease caused by severe acute respiratory syndrome coronavirus 2 (SARS-CoV-2) that had infected more than 16 million individuals and caused more than 656 000 deaths worldwide. A safe and effective vaccine against COVID-19 is urgently needed **[1]**. SARS-CoV-2, a member of the Betacoronavirus genus, is closely related to severe acute respiratory syndrome coronavirus (SARS-CoV) and several bat coronaviruses **[2, 3, 4]**. Compared to SARS-CoV and Middle East re-piratory coronavirus (MERS-CoV), SARS-CoV-2 appears to undergo more rapid transmission **[5, 6]** leading to the urgent demand for a vaccine. There are currently more than 160 COVID-19 candidate vaccines in development worldwide, and 25 are in different phases of clinical trials using different platforms **[7]**. Several vaccines, such as a recombinant adenovirus type-5 (Ad5)–vectored vaccine, a chimpanzee adenovirus-vectored vaccine (ChAdOx1 nCoV-19), and 2 mRNA vaccines, have been published or made available on preprint servers **[8**,**9]**. Inactivated vaccines have been widely used for the prevention of emerging respiratory diseases for decades, **[10]** and the relatively high speed of the development of this kind of vaccine makes it a promising strategy for COVID-19 vaccine development. Moreover, results from preclinical studies of 2 inactivated COVID-19 vaccines have shown that the vaccines could protect against SARS-CoV-2 with varying efficacy.

In spite of all the efforts conducted to fight against COVID-19, limited clinical and laboratory achievements have been reported so far. Thus, successful experimental steps for isolating and characterization of SARS-CoV-2 from patients are crucial for vaccine development. To develop inactivated SARS-CoV-2 vaccine candidate; isolation of the virus from Egyptian patients followed by propagation, characterization and virus inactivation was conducted. In vitro neutralization was performed in mice as efficacy and immunogenicity were tested for preclinical evaluation of the inactivated vaccine candidate for further clinical trial approach.

## Materials and methods

### Sample collection

Nasopharyngeal and oropharyngeal swabs were collected in 5 ml viral transport media from six COVID-19 patients with age >45 years, whom were positively diagnosed using rRT-PCR. Swabs were transferred to Egypt Center for Research and Regenerative Medicine (ECRRM) BSL-3 laboratory and stored at 4°C for immediate virus isolation. The approval of the ethics institutional review board (IRB) of Ministry of Defense, written informed consent was obtained from the participants.

### Biosafety containment

Using the precautions that adhered to or exceeded the requirements by WHO **[11]**, all experiments with suspected samples, infected cells, and infected fluids were performed in ECRRM Biosafety Level 3 laboratory and were conducted under appropriate conditions.

### Virus isolation, propagation and identification

Virus isolation was applied on Vero cell line of kidney epithelial cells derived from the African green monkey (ATCC No. CCL-81) supplied from Veterinary Serum Vaccine Research Institute (VSVRI). Confluent monolayer of Vero cells were grown in Dulbecco’s modified minimum essential medium (DMEM) supplemented with penicillin (100 units/ml), streptomycin (100 mg/ml), 0.2% sodium bicarbonate and 10% fetal bovine serum (FBS). The prepared cell culture was infected with 1 ml of suspected SARS-COV2 samples for about 45-60 minutes, then maintenance medium (MEM supplemented with 2% fetal bovine calf serum) was added and followed by incubation at 37±1°C. The cells were examined twice daily for cytopathic effects (CPE) formed by inoculated virus. Three blind passages **[12]** followed by seven successive serial passages were obtained and tissue culture suspensions were collected for virus detection and quantification by Real-time PCR. Virus replication and isolation were confirmed through cytopathic effects, gene detection, and electron microscopy.

### Real-Time PCR detection

Total RNA was extracted using Viral RNA Extraction kit (Qiagen, CA) following manufacturer’s instructions. Extracted RNA concentration and purity were tested with NanoDrop spectrophotometer (Thermo Fisher Scientific, USA). One step Real-time RT-PCR was achieved using TaqPath™ COVID-19 CE-IVD RT-PCR Combo Kit (Thermofischer Scientific, USA) following manufacturer’s instructions. The reaction was incubated in Real-Time PCR ABI 7500 (Thermo Fisher Scientific, USA) at 50°C for 15 min for reverse transcriptase step, 95°C for 10 min, followed by 45 cycles of 95°C for 15 s and 60°C for 30 sec. Three primers/probes were used targeting ORF1ab, Nucleocapsid (N) and Spike (S) regions. Primers/probes specific for bacteriophage MS2 were used as a positive control. The cycle threshold value below 33 was stately to be positive. Result is valid when two of the three targeted genes and the MS2 showed positive results.

### Whole Genome sequencing

Extracted RNAs were quantified using Qubit RNA High Sensitivity Kit (Invitrogen, USA). Libraries were prepared using Ion AmpliSeq SARS-CoV-2 Kit (Thermo Scientific, USA) following the manufacturer’s Protocol. Clonal amplification of the libraries was done using the Ion-PI-Hi-Q Sequencing 200 Kit (Thermo Scientific, USA) PCR emulsions. Purified libraries were qualified and quantified by Agilent Bioanalyzer and Qubit 4 Flurometer (Thermo Scientific, USA). Libraries were sequenced on the Ion proton NGS platform (Thermo Scientific, USA). Virus sequence assembly was performed using The Ion Torrent package (v.5.12) followed by genome mapping using tmap program (v.512) against complete SRAS-CoV-2 genome sequences retrieved from the GISAID website. All of the strains were isolated from Vero cells, which have been certified by WHO for vaccine production. Vero cells monolayer, were infected via the swabs of patients to prevent possible mutations during viral culture and isolation.

### Transmission electron microscopy

Infected Vero cells were scraped from the flask, pelleted, and cell pellet was rinsed with 0.1M phosphate buffer (Sigma Aldrich, Germany). Cell suspensions collected from inoculated VERO Cell monolayers were first fixed with 2% Formaldehyde in phosphate buffered saline for 1h before ultracentrifugation (1h, 25,000 rpm), loading sample on carbon coating grid stained with 2% phosphor-tungestic acid for 30 seconds then examined. Bar:100nm.

### Virus titration

The 50% tissue culture infectious dose (TCID_50_) per ml was determined in Vero cell monolayers on 24 and 96-well plates.Serial dilutions of virus samples were incubated at 37°C for 4 days and subsequently examined for cytopathic effect (CPE) in infected cells. TCID_50_ assay was performed according to **[13]**. The infectious titer was calculated using an in-house method adapted by Spearman and Kärber and expressed in TCID50 units **[14]**.

### Virus inactivation and vaccine production

For vaccine preparation, the virus was propagated in Vero cells with a dilution of 1:100 (v:v) of the SARS-CoV-2 virus in serum-free medium. The cells were incubated at 37^0^C for 72h. At three days post infection, when the cytopathic effect (CPE) was visible, the virus was harvested by collecting cell supernatants. The infectious titer of the virus was determined using a 50% cell culture infectious dose as described above. Vaccine purification was then performed with low-speed centrifugation (1000 rpm) to clarify the cell harvest followed by filtration. SARS-CoV-2 was inactivated with β - propiolactone at 2-8°C for 24-32 hours **[15**,**16]**. The final bulk was prepared as a liquid formulation containing 55 μg, or 100 μg total protein with aluminium hydroxide (Alhydrogel® CRODA health care Corp.) as adjuvant (0·45 mg/mL) per 0·5 ml.

### Validation of the inactivation

Effective inactivation of the virus was validated by inoculations of vero monolayers in 75 cm2 flasks with ten ml of inactivated virus, and then cultured at 36 ± 1°C for 4 days. No CPE was observed for three passages, in addition, quantitative PCR (Q-PCR) performed at several time points during passage confirmed the absence of amplification of virus genomes **[17]**.

### Safety Test

The vaccine formulations at two different concentrations and the adjuvant have been evaluated for safety in groups of Swiss albino mice (n=10/group). Safety has been documented in repeat-dose toxicity studies in mice (female, 6-8weeks old) which were vaccinated intraperitoneally (i.p) with three doses (N+1) at 55 and 110 µg/dose of inactivated vaccine candidate without adjuvant on day 0, 7 and 14 **[18]**. In contrast, mice group was treated (i.p) with a single dose of alum. Hydroxide at the dose of 5mg of Al (OH)**3** /mouse, which equivalent about 25 −8 human doses that recommended (0.2 - 0.8mg). Another negative mice group was injected with phosphate buffered saline (PBS) as control group. All animals were observed for mortality during the experimental period. Animals were euthanized on day 21 and 28 and necropsied and organs were evaluated for macroscopic and microscopic findings. Evaluations for histopathology marked as to the vaccine/dosage to assess the extent of pathologic damage and the eosinophilic component of the inflammatory infiltrates.

### Immunization of mice

Immunization of mice Female mice (6–8weeksold) were obtained from (VSVRI), after sampling of pre-immune sera. Mice were immunized with two different doses of the candidate vaccine 55 and 110 µg, total protein, adjuvanted with 0.5mg aluminium hydroxide (alum) to each antigen concentration. Groups of n=10 mice were inoculated intramuscularly (i.m.) with 0.5 ml of each vaccine preparation; control groups received the same volume of buffered saline. A booster immunization was carried out fourteen days post immunization with the same formulation and dosage as primary inoculation. Sera were then drawn from all groups every week to quantitative evaluation of SARS-COV2 neutralizing antibodies in vaccinated mice **(19)**.

### SARS-CoV-2 challenge test

Three weeks post the booster dose, all mice of the groups that received vaccine doses of 55 and 100 µg and the control group were challenged. Prior to challenge, a blood sample was drawn for determination of neutralising antibody titres. For challenge, mice were anaesthetised with isofluran and inoculated intranasally with with 60 µl of SARS-CoV-2 virus (10^6^ TCID_50_) according to animal care and use guidelines in an approved animal BSL-3 laboratory **[20]**. The isolated SARS-CoV2 that has been propagated five times on serum protein free Vero SF cells was used for homologous challenge. On day 3 and 7 post challenge, mice were euthanised, before lung and trachea were removed and frozen at −80°C. Tissue samples were thawed and homogenised in 1ml of Vero cell culture medium supplemented with antibiotics for titration in the TCID50 assay **[21]**.

### Histopathological analysis

The lung tissues of challenged mice were immediately fixed in 10% buffered formalin and embedded in paraffin wax. Histopathological changes caused by isolated SARS-CoV-2 virus infection were examined by H&E staining and viewed under the light microscope as described previously **[22]**.

### Determination of neutralizing antibody titers

Serum samples were heat-inactivated for 30 minutes at 56°C and serially diluted with cell culture medium in twofold dilution. The serum dilutions were mixed at a ratio of 1:1 with a SARS-CoV-2 virus stock suspension adjusted to 100 TCID_50_/ml, incubated for 1h at 37°C in a humidified atmosphere with 5% CO2 and transferred (eight replicates per dilution) to a 96-well tissue culture plate seeded with a Vero cells. The plates were incubated for 5 days at 37°C in a CO 2-incubator, before the cultures were inspected under a light microscope for the presence of a cytopathic effect (CPE) caused by SARS-CoV2, i.e. cell rounding and detachment. Neutralizing antibody titers were expressed as the reciprocal of the last dilution of serum that completely inhibited virus-induced CPE.

### Statistical Analysis

Neutralizing antibody titers, lung virus titers, histopathologic lesion score and eosinophilic infiltration scores were averaged for each group of mice. Comparisons were conducted using parametric and nonparametric statistics as indicated

## Results

### Isolation and propagation SARS-COV2 virus for vaccine candidate development

A primary virus seed was generated from a human isolate by ten passages on vero cells. Distinct CPE in cells monolayer infected by SARS-CoV-2 was detected following an incubation period of 2–3days post cells infection **(Fig. 1)** indicated that the virus grow well on Vero cells, so it was reflected that this was doubtless the optimal cell matrix for rapid vaccine development. This primary seed was further amplified to generate a seed virus bank, a working virus bank and a production virus bank. The production virus was then used to infect serum protein free vero cell cultures resulted in generation of high viral titres (10^7.5^ TCID_50_/ml).

**Fig. 1:**
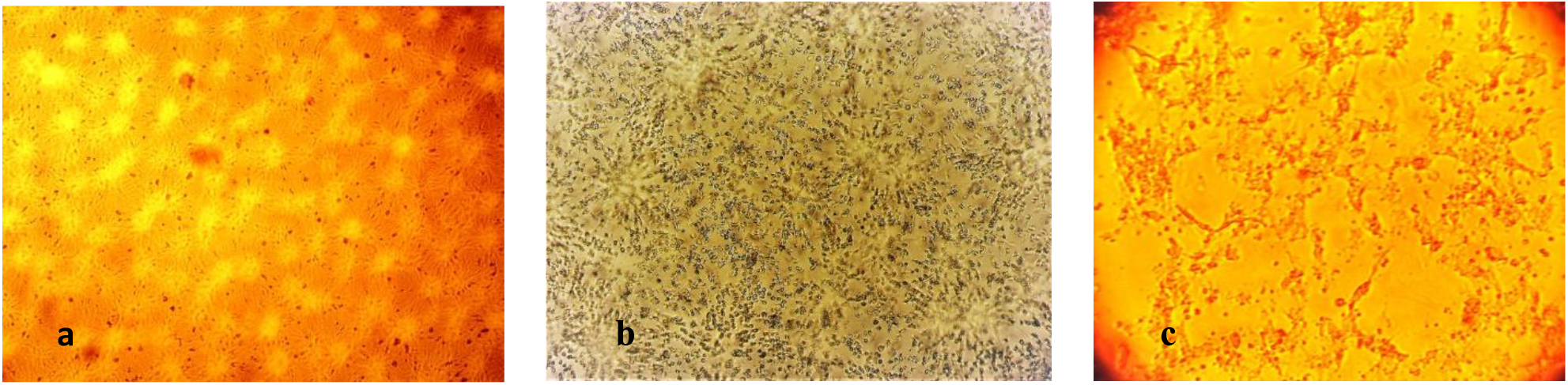
Normal Vero cells monolayer (a). Vero cell monolayer infected by SARS-CoV-2 48 hours post inoculation (b). Marked detachment of cells 72 hours post cell cultures infection (c).

### Determination of SARS-COV2 Virus Titer

Titration of the virus isolate revealed gradual increase in the virus titer through the successive passages **(Fig. 2)**. The virus titer was 5 log10 TCID50 at the 4th passage and reached 6.5 log10 TCID50 by the 8th passage. At the final passage, the virus titer reached 7.5 log10 TCID50.

**Fig. 2:**
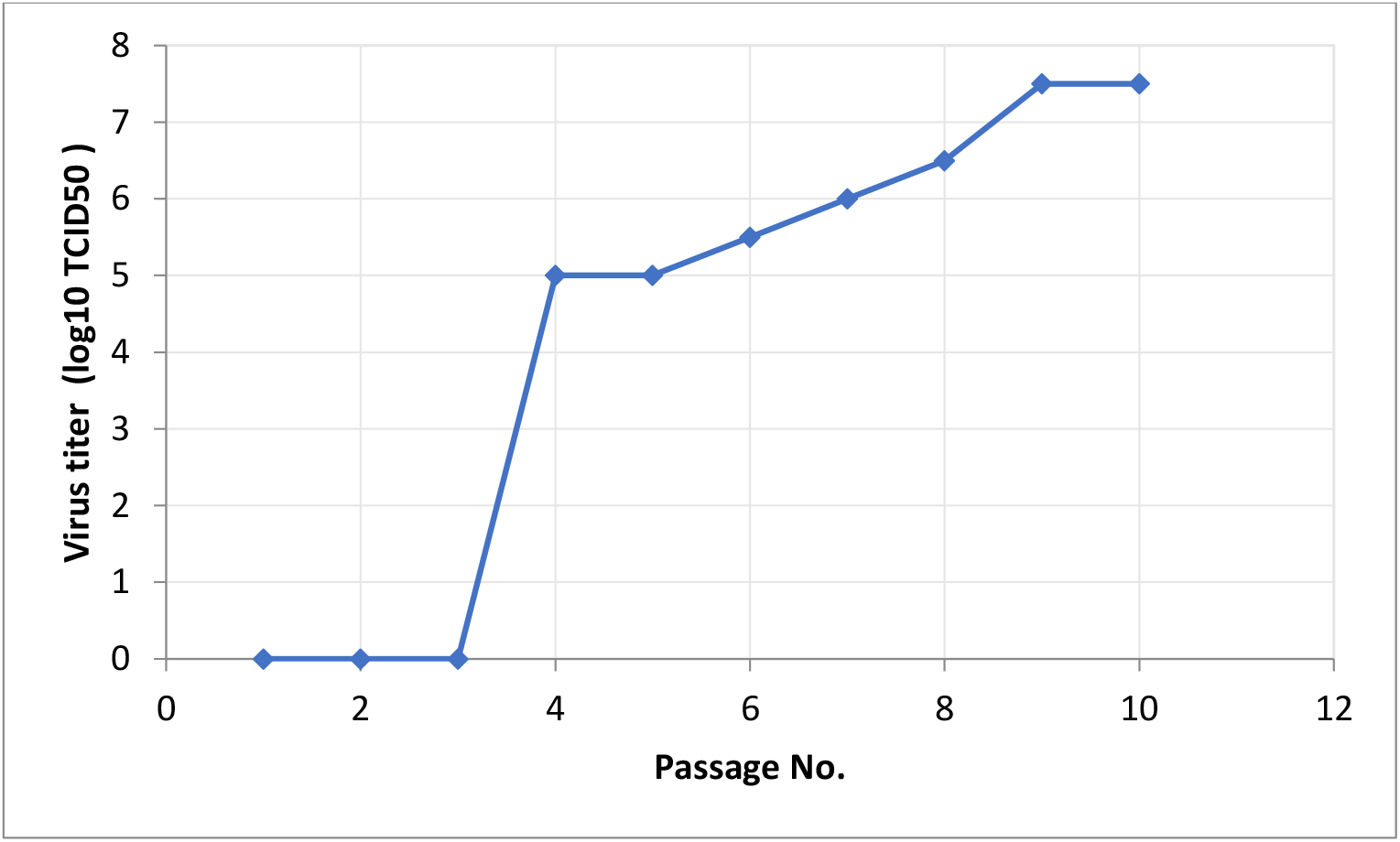
Titer of isolated SARS-COV2 in Vero cell culture.

### Whole Genome sequencing

complete viral genome sequences of the isolate’s SARS-CoV-2/human/EGY/Egy-SERVAC/2020 with accession numbers; MT981440; MT981439; MT981441; MT974071; MT974069 and MW250352 at GenBank revealed that the virus was most closely related (99.5% nucleotide similarity) to USA/VA-CDC-6377/2020 strain (MT325612.1) and USA/FL-BPHL-0259/2020 strain (MT704077.1).

### SARS-COV-2 inactivation and mouse safety test

The harvested virus containing supernatant was inactivated by β - propiolactone treatment at 2-8°C for 24. Inactivation was confirmed by three passages into Vero cells. No CPE was observed in the inactivated virus infected cell monolayers. An electron micrograph of the purified inactivated virus showed virion particles belonging to the Coronaviridae Family were observed. Negatively stained electron microscopy image visualized oval viral particles with spikes having diameters of approximately 100 nm (**Fig. 3**) confirmed that virus particles being demonstrated to present well defined spikes on the virus membrane.

**Fig.3:**
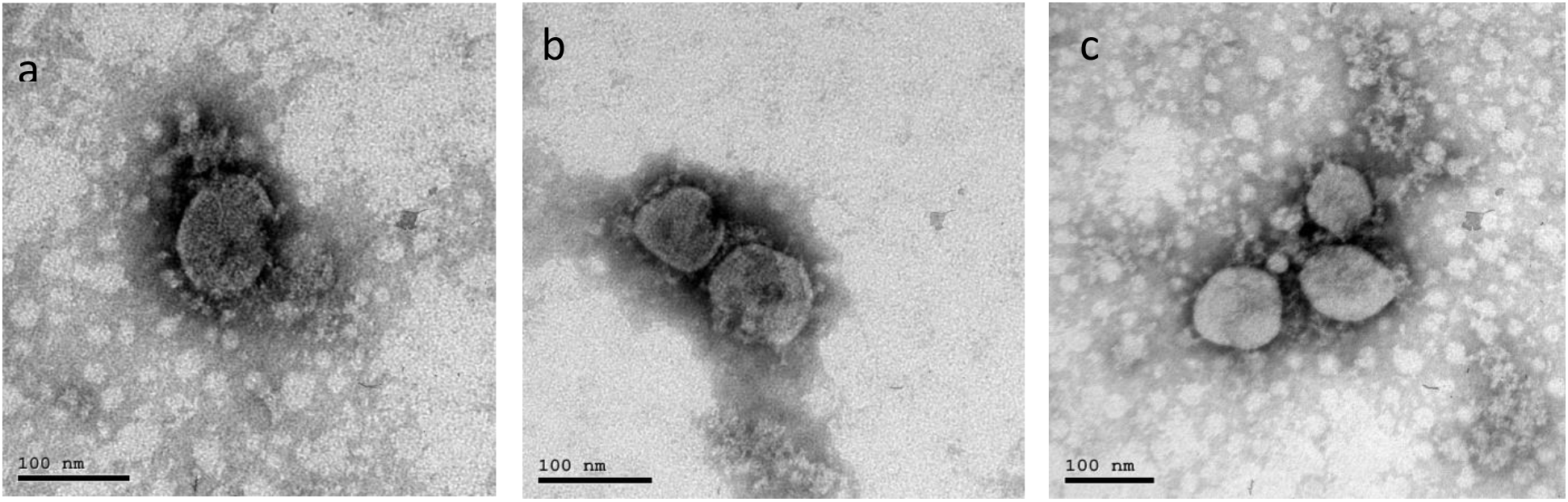
Virion Particle belonging to the Coronaviridae Family observed by electron microscopy (a).Electron micrograph (187,000-fold magnification) of purified inactivated SARS-CoV-2 candidate vaccine after staining with uranylacetate. Spikes formed by S protein project from the viral surface (b&c)

No mortality or morbidity was observed in mice inoculated intraperitoneally with repeated doses of the vaccine and adjuvant. Moreover the experiment revealed that the inactivation steps were independently capable of inactivating this titre with a large margin of safety **(Fig. 4&5)**.

**Fig.4:**
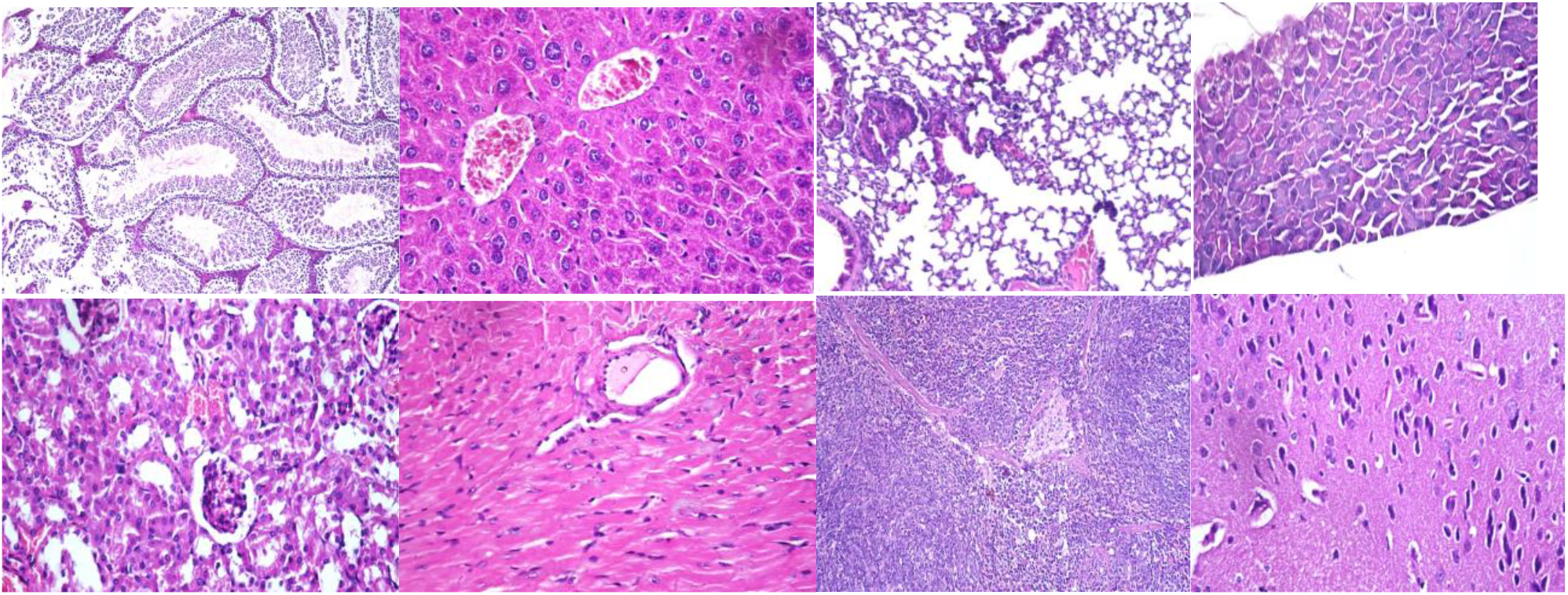
Normal organs tissue of mice vaccinated with three doses (N+1) of candidate inactivated SARS-COV2 vaccine **(H&E X 200)**.

**Fig.5:**
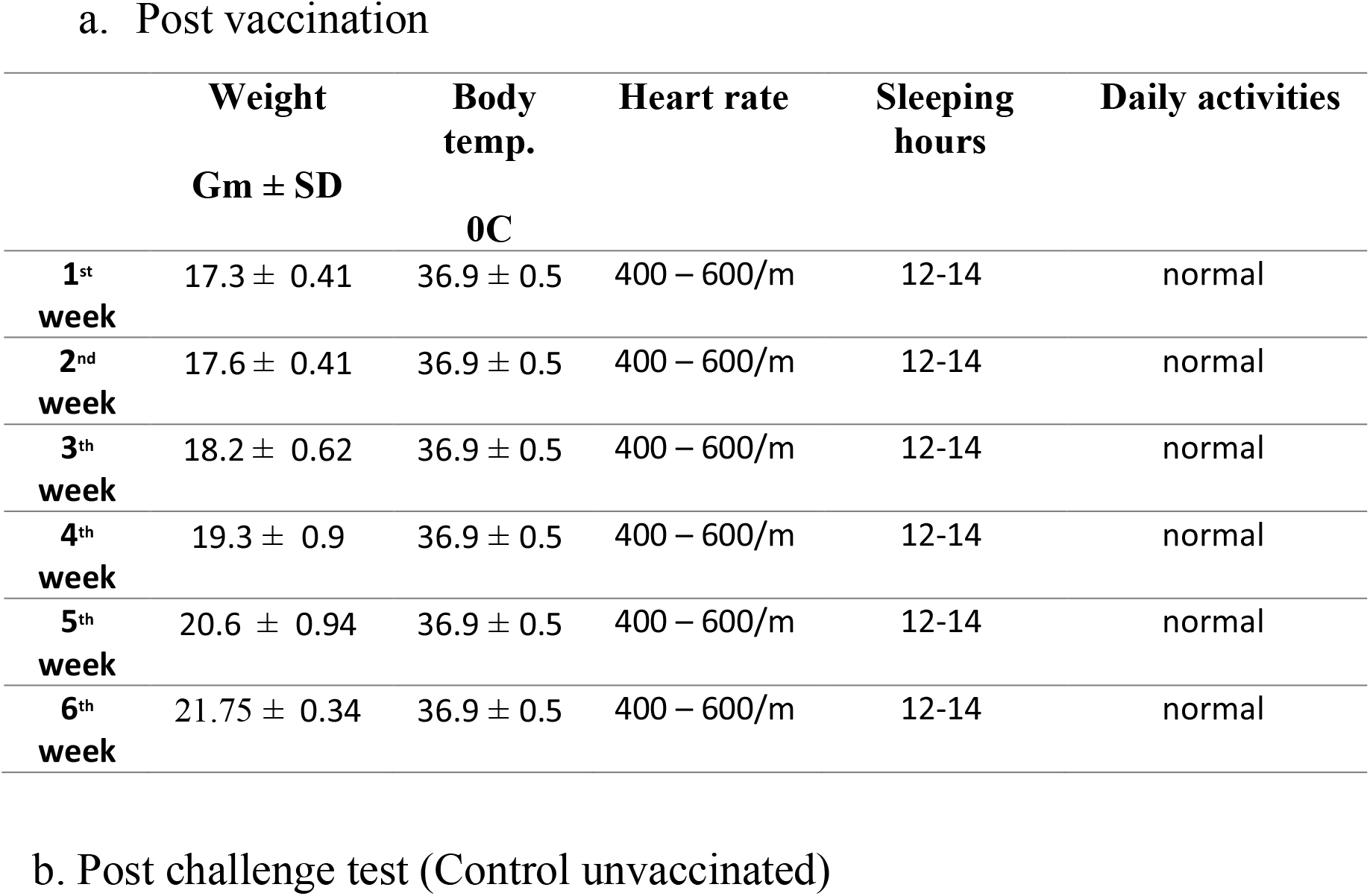
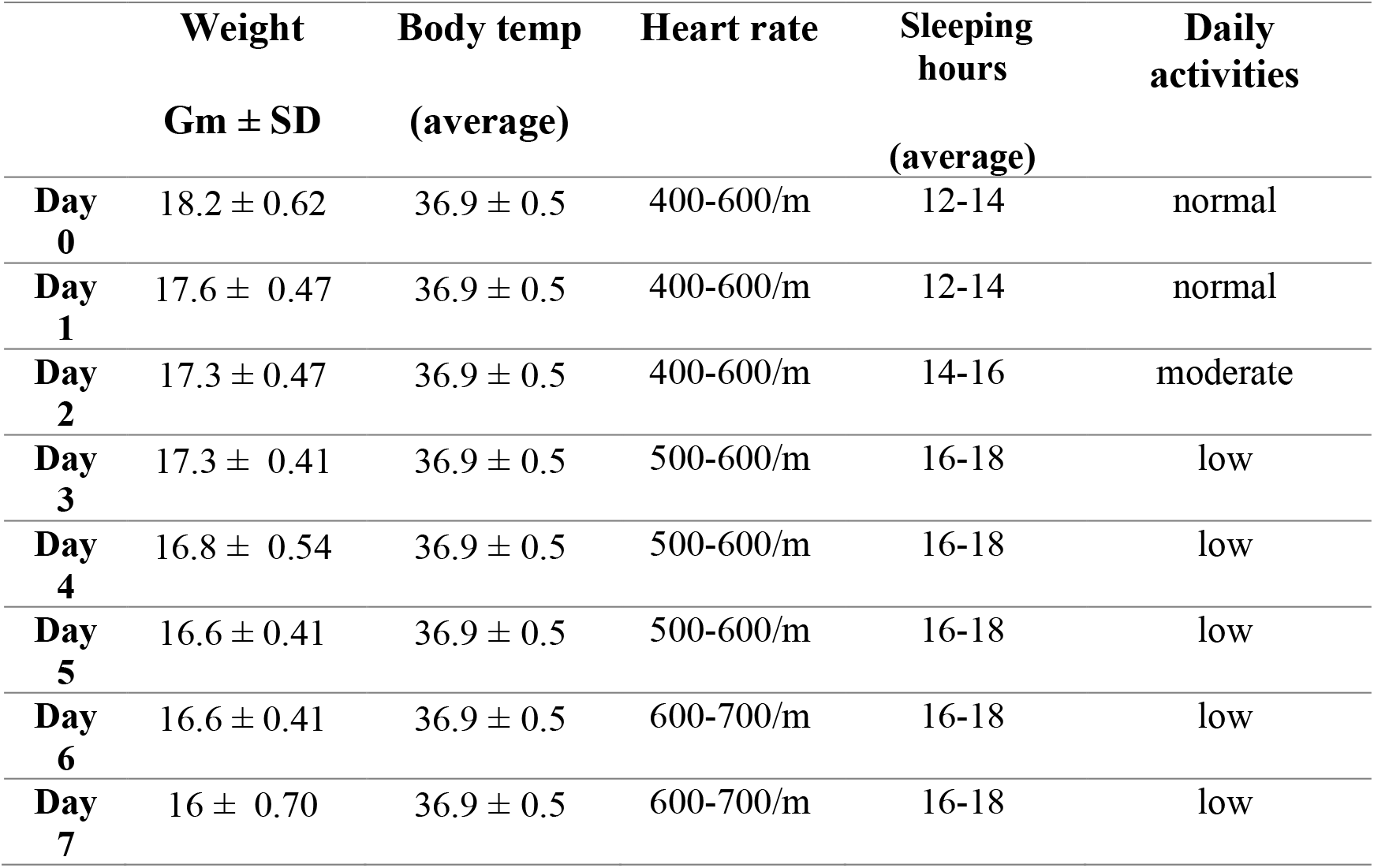
clinical signs observation post vaccination (a) and challenge test (b)

### Immunogenicity in mice

The immunogenicity of the candidate vaccine was initially investigated in dose-finding and adjuvant studies performed in mice. Neutralizing antibody titre determined for each group of animals is presented in **Fig. (5)**. No dosage effect was noted as the mean neutralizing antibody titers of the two doses of inactivated vaccine groups were non-significant different (p≥0.05). These data demonstrate that the candidate vaccine is highly immunogenic in mice. Following a single immunization with 55 or 100µg, SARS-COV2 specific antibody titres up to 1:213 were detected two weeks post first vaccination. Following the booster immunization, titres were substantially increased up to 1:2560 with the 100 µg dosage adjuvanted with aluminium hydroxide. High titre neutralising antibodies (approximately 1:1707) were also measured for the 100µg one week after the challenge test with 10^6^ TCID_50_ of SARS-CoV-2. All of the vaccinated animals had higher serum neutralizing antibody titers than non-vaccinated challenged mice group

### Efficacy in mice

Mice were monitored daily post challenge for morbidity (weight) and mortality. No clinical signs were observed in vaccinated groups compared with unchallenged control group. Unvaccinated group showed rough hair with arched back 3 days post challenge. Candidate vaccine was highly efficacious in mice at the two dosage levels. The challenge virus replicated to titers 10^2.5^ TCID_50_ /ml and 10^5.4^ TCID_50_ in vero cell culture infected with homogenized lung tissues from mice at 3 and 7 days post infection, respectively. Replication of the challenge virus was not detected by Real-time PCR the lungs tissues of mice that received the vaccine and by absence of CPE in cell culture monolayer (**Fig. 6**).

**Fig.6.**
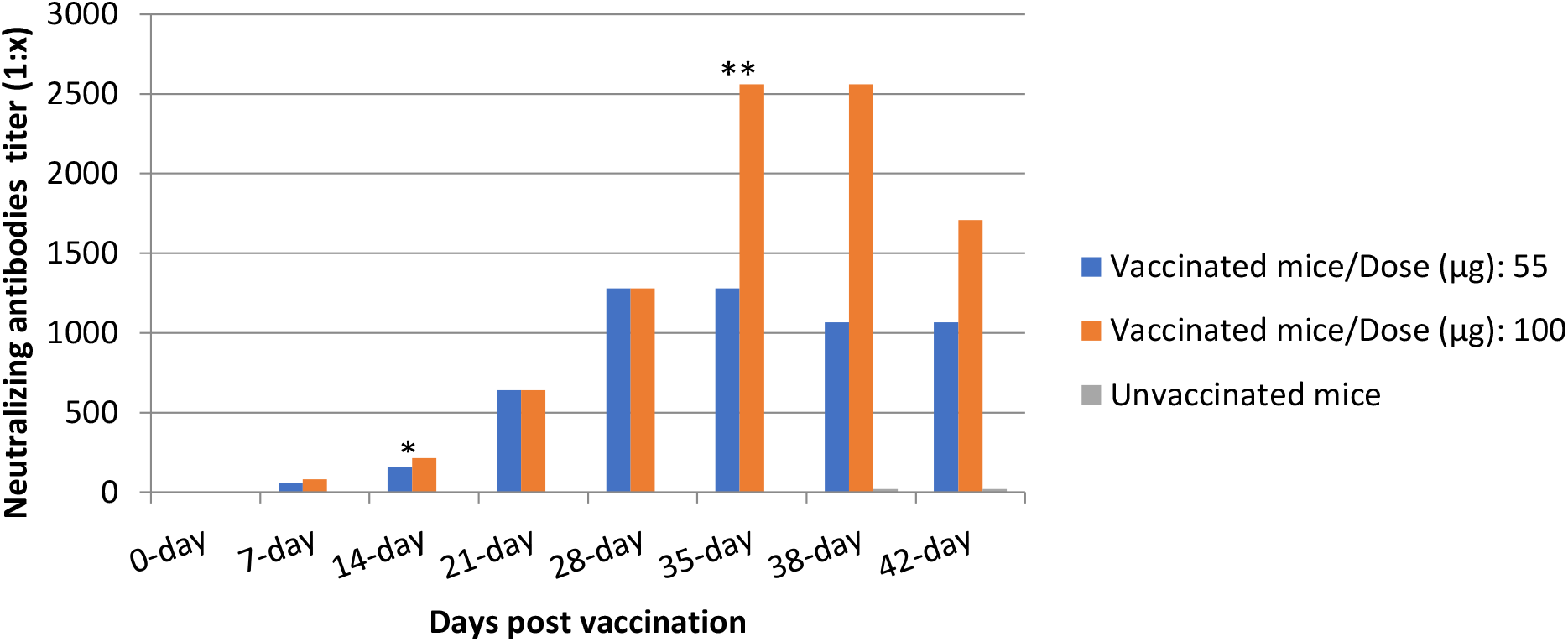
Mean neutralizing-antibody responses to the SARS-COV2 inactivated vaccine in mice. * Mice were re-immunized with booster dose **Mice were intranasally challenged with 10^6^ TCID_50_ of SARS-CoV2.

### Histological evidence of protective efficacy post challenge

Histopathological findings did not vary within either low dose (55µg) or high dose (100µg) vaccinated groups among tissues collected post challenge (Fig. 8). In addition to the reduction in viral titers detected in the lungs, histopathological findings in the lungs of immunized mice indicate that the candidate inactivated vaccine produced protection from SARS-CoV-2 after 3 and 7 days post challenge. It is meaning that vaccinated groups that had detectable levels of serum neutralizing antibodies to SARS-CoV-2 at the time of challenge were protected from severe lung lesions. The unvaccinated control animals that lacked detectable levels of SARS-CoV-2 neutralizing antibodies, had severe lung lesions, mice lung at 3&7days post challenge test showing diffuse thickening in the interstitial tissue, congested peri-alveolar blood capillaries and lymphocytic cells infiltrations.

**Fig.7.**
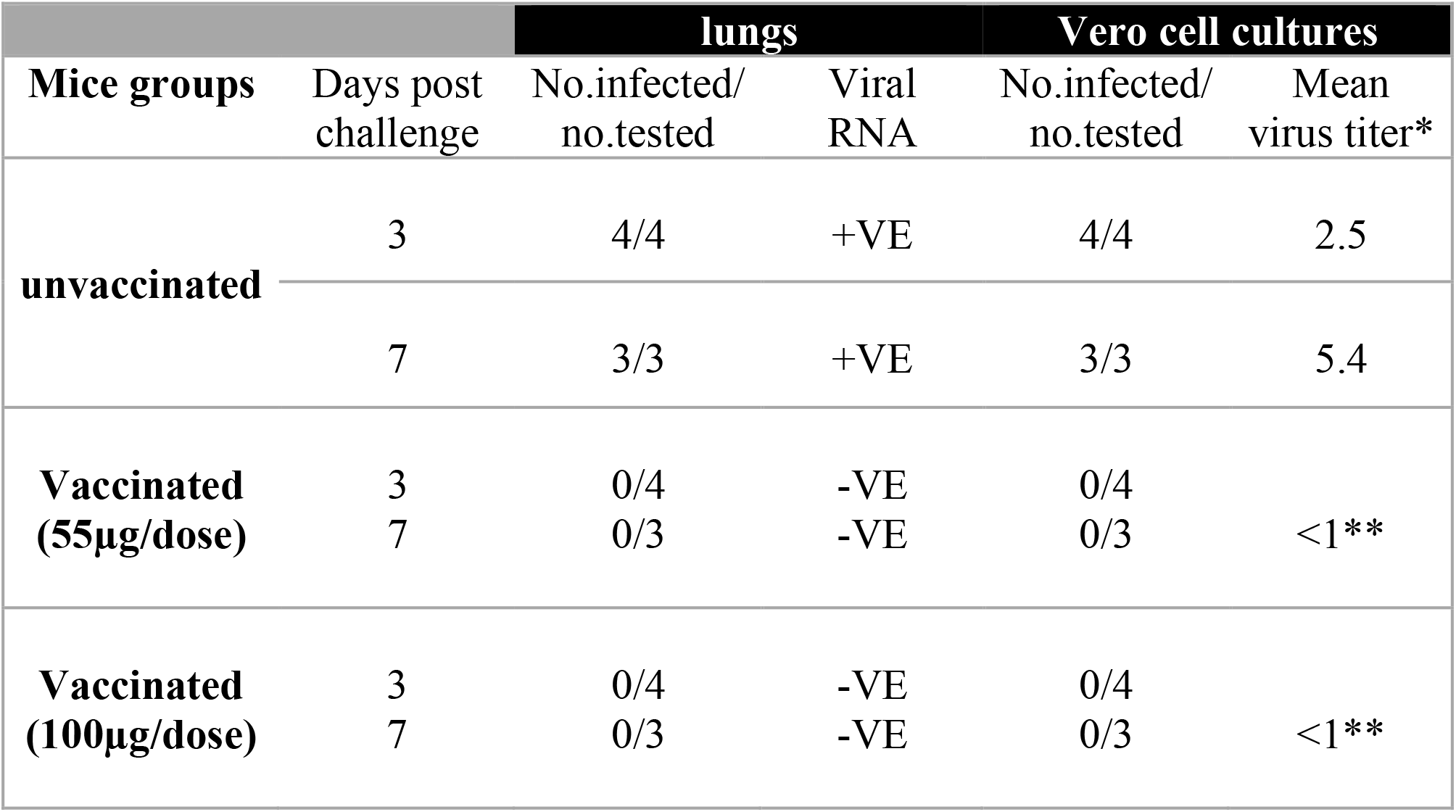
Virus replication in lung tissue upon challenge. *Virus titers are expressed as log_10_ TCID_50_/ ml of homogenized lung tissue. ** Virus not detected by absence of CPE in infected vero cell monolayer cultures.

**Fig.8:**
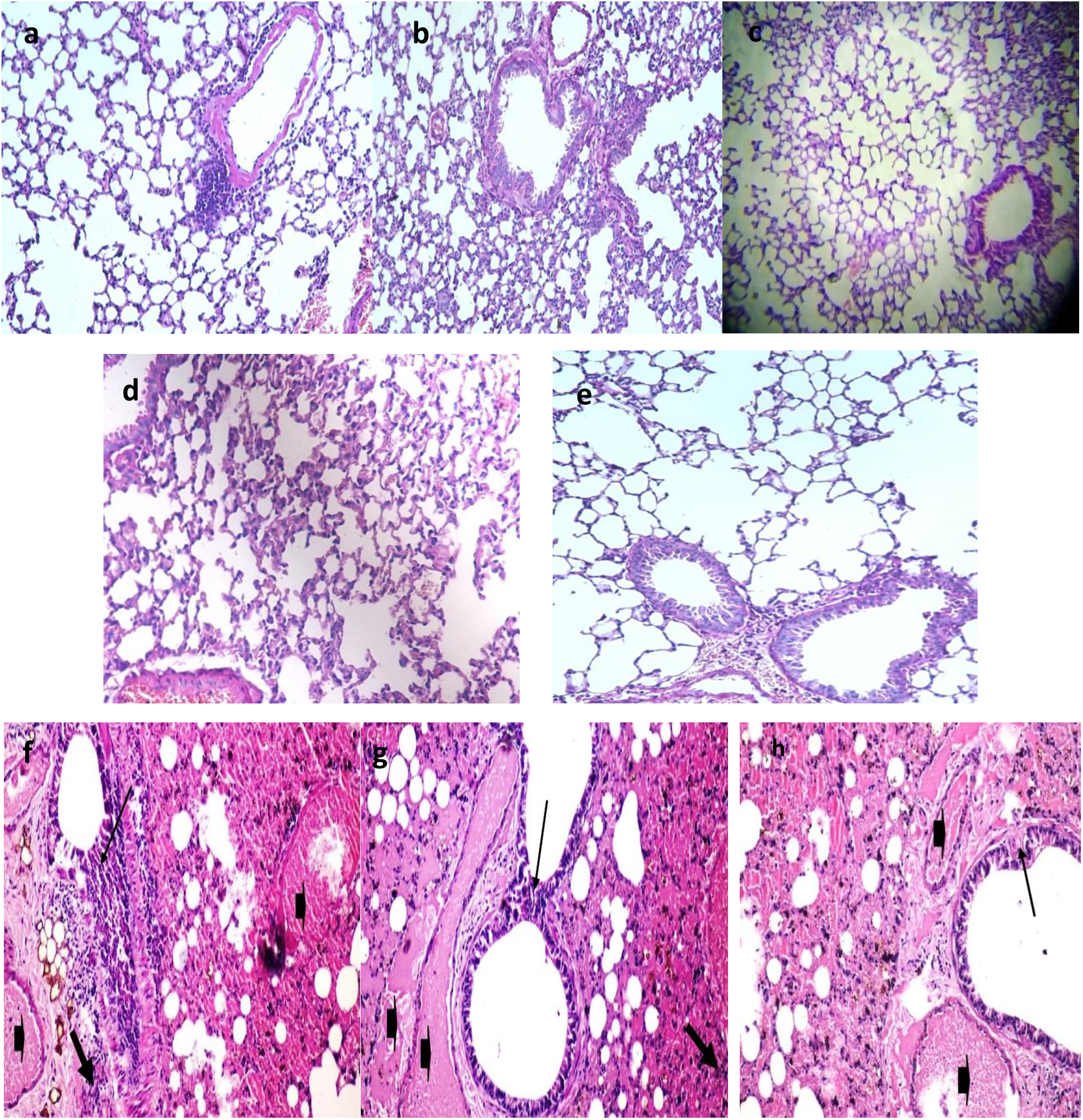
Normal lung tissue (a). Vaccinated mice with 55 (b) and 100µg (c) at 3 days and 7 days (d&e) post SARS-COV-2 challenge showing normal lung tissues. Unvaccinated mice lung at 3&&7days post challenge test showing diffuse thickening in the interstitial tissue, congestion peri-alveolar blood capillaries and lymphocytic cells infiltrations (f, g & h).

## Discussion

Novel COVID-19 has shown a rapid spread since December 2019 causing a huge outbreak in China **[23]**. Isolating and studying the causative virus is crucial for developing diagnostic tools, therapeutics, and effective vaccine. The development of vaccines with high immunogenicity and safety is of paramount for controlling the pandemic COVID-19 and prevent further infection spread. A number of different strategies have previously been reported for the development of experimental and candidate human SARS vaccines. These include inactivated whole virus vaccine **[24]** using large-scale serum protein free vero cell cultures was reported. The whole-genome sequence of isolated SARS-CoV-2 strain was closely related to most available sequences, representing to some extent circulating SARS-CoV-2 populations. As well as the development of SARS-CoV-2 vaccine was based on adaption of this isolated virus, to establish optimal conditions for growth, inactivation and purification of the inactivated virus. Inactivated vaccine derived from isolated SARS-CoV-2 strain which closely related to Wuhan/WIV04/2019 was reported **[19]**. The SARS-CoV-2 had been reported to grow effectively on Vero cells **[25, 26]**, considering that this was probably the optimal cell matrix for rapid vaccine development. Propagation of isolated virus on vero cell monolayers yield a titer of 7.5 log10 post ten passages with distinct CPE within 72 hr post infection. This was in agreement with **Gao et al. [19]** who discussed that vero cell cultures showed SARS-COV 2 replication efficiently and reached a peak titer of 6 to 7 log10 TCID50/ ml by 3 or 4 days pi. Previous findings recorded the SARS-CoV cytopathic effects with monolayers of Vero cells three days post the blind passages **[7]**. In contrast, no specific cytopathic effects were observed in the Vero E6 cells until 6 days after inoculation as reported by **Zhu et al. [18]**. Most processes for inactivated whole virus vaccines have utilized β-Propiolactone as an inactivating agent. Our study report development of an inactivated SARS-CoV-2 vaccine **(Egy-CoVac)** showed intact, oval-shaped particles with diameters of 90 to 150 nm, which were embellished with crown-like spikes, representing a pre-fusion state of the virus **(Fig. 3)**. These results are consistent with previous report for transmission electron microscopy analysis of stained samples demonstrated that the inactivated virion presented well defined spike structures on the virus particle with no apparent structural alterations resulting from the inactivation procedures **[24]**. In addition to, **Egy-CoVac** toxicity and safety evaluation showed no adverse or clinical signs in vaccinated <u>mice</u>. Inactivated whole virus vaccine would be most efficient in inducing neutralising antibodies, which are possibly critical in preventing SARS-CoV infection **[24]**. Since mice are a model of SARS-CoV infection but not disease so, Balb/C or Swiss Albino mice were used in evaluation of developed SARS-COV2 vaccine as previously reported **[27]**. It was reported that Whole Killed vaccine convened more protection against pulmonary SARS-CoV replication in mice lung tissue vaccinated with 50 µg inactivated virus in 0.2ml **[21]**. Our results show that the candidate vaccine formulations induced significantly elevated antigen neutralizing antibody responses in the vaccinated mice with low and high doses, protecting them against SARS-CoV-2 infection. These were in agreement with previous results of development inactivated SARS-COVI 2 vaccines, BBIBP-CorV and PiCoVacc from China and BBV152 whole virion inactivated vaccine **[19, 27, 28, 29]**. Alum.hydroxide gel which the most frequently used as vaccine adjuvant with an extensive safety record desired to have a COVID-19 vaccine that can generate both humoral and cell-mediated immune responses. The response generated from alum is primarily Th2-biased with the induction of strong humoral responses via neutralizing antibodies **[30]**. Although previous studies in mice have shown that low levels of neutralizing antibodies are sufficient to prevent detectable viral replication following challenge **[31, 32]**. In our study the viral RNA was detected by Real-time PCR in mice lung tissues harvested from the unvaccinated mice post infection as well as the mean SARS-CoV-2 titer of homogenized lung tissues in vero cell culture from the control group were 10^2.5^ TCID_50_ /ml and 10^5.4^ TCID_50_ at 3 and 7 days post infection respectively. Our data also demonstrate complete protection against SARS-CoV-2 challenge by inhibition virus replication in lung tissue post challenge test. These results concluded that this vaccine will not cause antibody-dependent enhancement (ADE) as all the data obtained in this trial support the safety and immunogenicity [the ability to incite an immune response] of this inactivated vaccine. As well as replication of the challenge virus was not detected in the lungs tissues of mice that received the vaccine. As antibody-dependent enhancement (ADE) of virus infection is a phenomenon in which virus-specific antibodies enhance the entry of virus, and this occur in case of nonsufficient antibodies that bind to the surface proteins but do not inactivate the virus

These data were confirmed by the absence of histopathological findings in the lungs of vaccinated mice groups. Same finding was recorded in mice experimentally infected with SARS-COV and SARS-COV2 virus as virus challenge was successfully established in animal models **[19]**. It was reported that a combination of high neutralizing antibody titers elicited against inactivated antigen alone and the presence intact spike protein on the surface of the virus confirms that the antigen is in the right formula and can itself may act as a Th1 inducer with its surface glycoproteins, intracellular viral proteins **[28]**. Therefore, Inactivated SARS-CoV-2 vaccine (**Egy-CoVac)** described here provided potential solution to fight against COVID-19 pandemic and has desirable properties that support further development and studies for clinical trials.

## Declaration of Competing Interest

The authors have declared no conflict of interest

## Conflict of Interest

The authors have declared no conflict of interest

## Compliance with Ethics Requirements

All Institutional and National Guidelines for the care and use of animals were followed.

## Highlights

Trial for an inactivated whole SARS-CoV-2 vaccine candidate development

Preliminary evaluation of the vaccine immunogenicty induces high levels of neutralizing antibodies titers in mice model, regardless dose concentration, that produce complete protection from wild virus challenge.

## Notes

### Competing Interest Statement

The authors have declared no competing interest.

